# PKS-A Clade of Oil Palm Might Play Role During Defense Against *Ganoderma boninense* Infection

**DOI:** 10.1101/2020.08.23.263871

**Authors:** Zulfikar Achmad Tanjung, Redi Aditama, Condro Utomo, Tony Liwang, Reno Tryono

## Abstract

Polyketide synthase (PKS) is an essential catalyzing enzyme in the polyketide (PK) biosynthesis pathway of bacteria, fungi and plants which have diverse beneficial functions such as antibiotic and antiparasitic. This study was aimed to identify specific plant type III PKSs in the African oil palm, *Elaeis guineensis*, and predict its biosynthesized metabolites as plant defense compounds against the most threatening fungal pathogen, *Ganoderma boninense* that causing the basal stem rot disease. We used the oil palm protein database to detect the presence of type III PKS domains using the HMMER version V3.1b2. An artificial inoculation was made on oil palm root tissues and RNA sequencing was performed to obtain the transcriptome profile after 7 days exposure to *G. boninense*. Among 40,421 proteins, we identified 38 of which containing type III PKS domains. Signal peptide signature motifs were absence in all PKSs suggesting their intracellular functions during the polyketide biosynthesis. A molecular phylogeny analysis reflected the relationships among these PKSs that clustered into PKS-A, -B and -C clades. Most of the PKS-A members were up-regulated after *G. boninense* infection, indicating their essential role in the biosynthesis of PK products which might needed for defense.

---

The Polyketide (PK) is a class of major secondary metabolite product, beside terpenoids/steroids, alkaloids, and non-ribosomal polypeptides, produced by bacteria, fungi and plants (Lussier *et al*. 2012; Pickens *et al*. 2011). It is known to have diverse and important biological functions including antibiotic and antiparasitic pytoalexin (Jez *et al*. 2000; 2001; Staunton and Weissman, 2001). Rapamycin (polyene immunosuppresent), erythromycin A (macrolide antibiotic), lovastatin (anti-cholesterol) and amphotericin B (polyene antifungal) are the examples of wide known PK products (Staunton and Weissman, 2001). PK is built from acetyl coenzyme A and malonyl coenzyme A backbone and iteratively synthesized carbon chain elongation by a series of intermolecular Claisen condensation. The lengthening and cyclization of polyketide chain are controlled by an essential enzyme named as polyketide synthases (PKSs) that catalyze the condensation reaction (Flores-Sanchez and Verpoorte, 2009).

According to its architectural configuration, there are three types of PKSs (Shen, 2003): Type I is a group of multifunctional enzymes that are harbored to a set of distinct module with non-iterative function, responsible for the catalysis of one cycle of polyketide chain elongation as exemplified by the 6-deoxyerythromycin B synthase (DEBS) for the biosynthesis of reduced polyketides (i.e. macrolides, polyethers and polyene) such as erythromycin A; type II is a monofunctional enzyme group with a single active site for each individual enzyme; and type III acts directly on acyl-CoA substrates which lack acyl carrier protein (ACP) (Pickens *et al*. 2011). Specifically, type III PKS enzymes involve in the plant secondary metabolite biosynthesis of compounds which responsible for pigmentation, fertility, and defence against pathogens or phytoalexins (Lussier *et al*. 2012; Schroeder, 1997a). It has been shown in sorghum that the activity level of chalcone synthase, a member of type III PKS, was significantly increased after inoculated with a common cereals pathogen, *Colletotrichum graminicola* (Lue *et al*. 1989; Nicholson *et al*. 1987). Similarly, the transcript accumulation of barley chalcone synthase in the inoculated leaves with *Bulmeria graminis* f.sp. *hordei* was abundantly accumulated, indicating its role in plant defense against pathogen (Christensen *et al*. 1998). Nowadays, plant PKSs exploitations are limited to those species with seasonal life cyle, despite there are many other annual trees with unknown PKS function to be revealed such as the oil palm.

The African oil palm, *Elaeis guineensis*, is well known as the most productive oil-bearing crop in the world. This species is intensively cultivated in South East Asia mainly Indonesia and Malaysia. Both countries contribute 85% of the world’s palm oil production (Yan, 2017). The genome was sequenced with a total of 1.8-gigabase (Gb) with 34,802 genes that distributed in 16 chromosomes, and it opens for public in the genome databank (Singh *et al*. 2013). In spite of the successful story of oil palm, the crop is facing the most threatening Basidiomycete fungal pathogen, *Ganoderma boninense*, that causes a devastate disease called the basal stem rot (Rees *et al*. 2009). It caused more than 50% yield losses and economic impact up to 500 million USD a year (Cooper *et al*. 2011; Hushiarian *et al*. 2013). Up to now, resistant variety is considered as a plausible approach to impede the disease development and thereof studies to apprehend plant defense mechanism against this pathogen remain open (Sahebi *et al*. 2015). This study was aimed to identify type III PKSs in oil palm and co-analyzed their transcriptional level during exertion period to the pathogen. According to our knowledge, this study is a first report exploring oil palm genome sequence to reveal PKSs and predict their role as plant defense compounds.

## MATERIALS AND METHODS

### Protein database mining

The *E. guineensis* protein database was downloaded from http://www.ncbi.org_under_accession_no_GCF_000442705.1CG5. The database was stored in our local server for further analysis. In parallel, we downloaded the FAE1_CUT1_RPPA (PF08392), chalcone and stilbene synthases N- (PF00195) and C-terminal domains (PF02797) from the Pfam database at http://pfam.xfam.org. These domains were described as type III PKSs in plants. HMMER version V3.1b2 (http://www.hmmer.org) was used to detect whether any of these domains is present in *E. guineensis* proteins. Each predicted PKS was re-analyzed to Pfam 31.0 database through “sequence search” tool to identify the PKS domains positions in the respective amino acid sequence. In each confirmed PKS, the domains were illustrated as colored-horizontal bars using an open source software Illustrator of Biological Sequences (IBS) version 1.0 (Liu *et al*. 2015).

### Phyllogeny analysis

Molecular phylogeny was built using the software DNASTAR Lasergene MegAlign version 7.0.0. Nine putative PKSs of date palm (*Phoenix dactylifera*) and five *Arabidopsis thaliana* at the NCBI database were included. *Phoenix dactylifera* represents the closest genetic relationship to *E. guineensis* among other palmae members while *A. thaliana* represents a common plant model as an out-group. The amino acid sequences for both species could be accessed under the following accession numbers, respectively: XM_008776974, XM_008780531, XM_008781792, XM_008782591, XM_008798885, XM_008809970, XM_008810587, XM_008810589, XM_008814523, NM_100085, NM_116221, NM_119651, NM_121396, and NM_001342307. For all PKSs, alignments were created by implementing the ClustalW algorithm and phylogenetic tree was inferred using the Neighbor-Joining method. The confidence of the resulting tree was estimated with 1,000 bootstrap iterations.

### Signal peptide prediction

All PKSs were analyzed for the presence of signal peptide motif using an online software the SignalP version 4.1 to predict whether the enzyme being secreted outside the cells or else (Petersen *et al*. 2011). Each protein sequence of PKS was modeled using the SWISS-MODEL software (http://www.swissmodel.expasy.org) prior to protein 3 dimensional (3D) construction (Biasini *et al*. 2014). The 3D visualization of a representative PKS was done using the software VMD 1.9.2 to obtain particular protein conformation structure in each PKS clade (http://ks.uiuc.edu/development/Download/download.cgi?packageName=VMD).

### Transcriptional analysis

The *G. boninense* G3 strain was used to challenge two months old oil palm ramets *in vitro*. A long cultured tube (150 x 25 mm) was filled with one gram of sterile rubber wood sawdust (RWS) and 2 ml of pre-solidified potato dextroxe agar (PDA) medium. This was autoclaved at 121°C, 15 psi for 15 minutes using Hirayama HVA-110 (Hirayama Manufacturing, Tokyo, Japan) and incubated at 25°C for three days to show if there was no contaminant occurred because of sterilization failure. A seven-days old mycelia plug of G3 culture (Ø 50 mm) was transferred into the tube and incubated at 30°C for 21 days. As control, no fungal mycelium was added into the tube. Subsequently, 7 ml of Murashige and Skoog (MS) medium with 10 g/l gelrite were added on top of the fungal culture. After three days, oil palm ramets were sub-merged into the MS medium until the root ends attached to the fungal culture. This was incubated at 28°C with 16 hours light from T8 Philips lamps 28 Watt (3,000-4,000 lux) and 50-60% relative humidity for 7 days post inoculation (dpi).

Ramet root tissues were harvested at 7 dpi from both infected and mock treatment as control. In each, there were four plants that used as biological repeats. Total RNA of each sample was extracted using the RNeasy^®^ Plant Mini Kit (Qiagen, Hilden, Germany) following manual instruction. RNA quality was analyzed using the QIAxcel system (Qiagen, Aldermaston, USA) and RNA quantity was measured using the spectrophotometry NanoDrop 2000C (Thermo Scientific, Massachusetts, USA).

RNA Sequencing Analysis was conducted using Illumina Technology (Illumina, San Diego, CA, USA). The FASTQ files contained paired-end short reads data were analyzed using Fastx toolkit for quality filter with minimum Phred value of 20. The filtered fragments were mapped to *Ganoderma sp. 1059 SSI* transcript model obtained from the Joint Genome Institute (JGI) (genome.jgi.doe.gov) under project id: 1078003 to exclude mRNAs belongs to *G. boninense*. Unmapped reads from previous process were mapped to *Elaeis guineensis* transcript obtained from NCBI Refseq under an accession GCF_000442705. All reads were mapped using Burrows-Wheeler Aligner (BWA) MEM which is suitable for Illumina short reads (Li and Durbin 2009). The transcripts were quantified from alignment files using eXpress software (bio.math.berkeley.edu/eXpress). The quantity of transcript was stated using transcript per million (tpm) terms. All transcripts were annotated against UniProtKB/Swiss-Prot database using blastx software with E-value restriction of 10^-5^. Blast result was used for GO terms analysis using Blast2GO software.

The relative expression was calculated by dividing each mRNA-ID transcription’s abundance of *Ganoderma*-treated with mock control transcriptomic databases. This was used to gain a hint whether any of the present PKSs play role as plant defense compound against *G. boninense*.

## RESULTS AND DISCUSSION

### Protein mining for PKS

The oil palm protein database in the NCBI databank stored 40,421 proteins that were used in this study. We attempted to identify PKSs through the presence of any of the following type III PKS domains i.e. FAE1_CUT1_RPPA (PF08392), chalcone and stilbene synthases N-(PF00195) and C-terminal domains (PF02797)) using HMMER program. Subsequently, we re-analyzed each predicted PKS against the Pfam 31.0 database through “sequence search” tool to identify the PKS domains positions in the respective amino acid sequence. This harbored 38 chalcone and stilbene synthase-like which known as specific type III PKSs in plants. Moreover, Pfam 31.0 identified any of the three characteristic PKS domains by their FAE1/type III polyketide synthase-like protein, N- and C-terminal of chalcone and stilbene synthases signature motif in all identified PKSs **Table 1**.

**Table 1.**
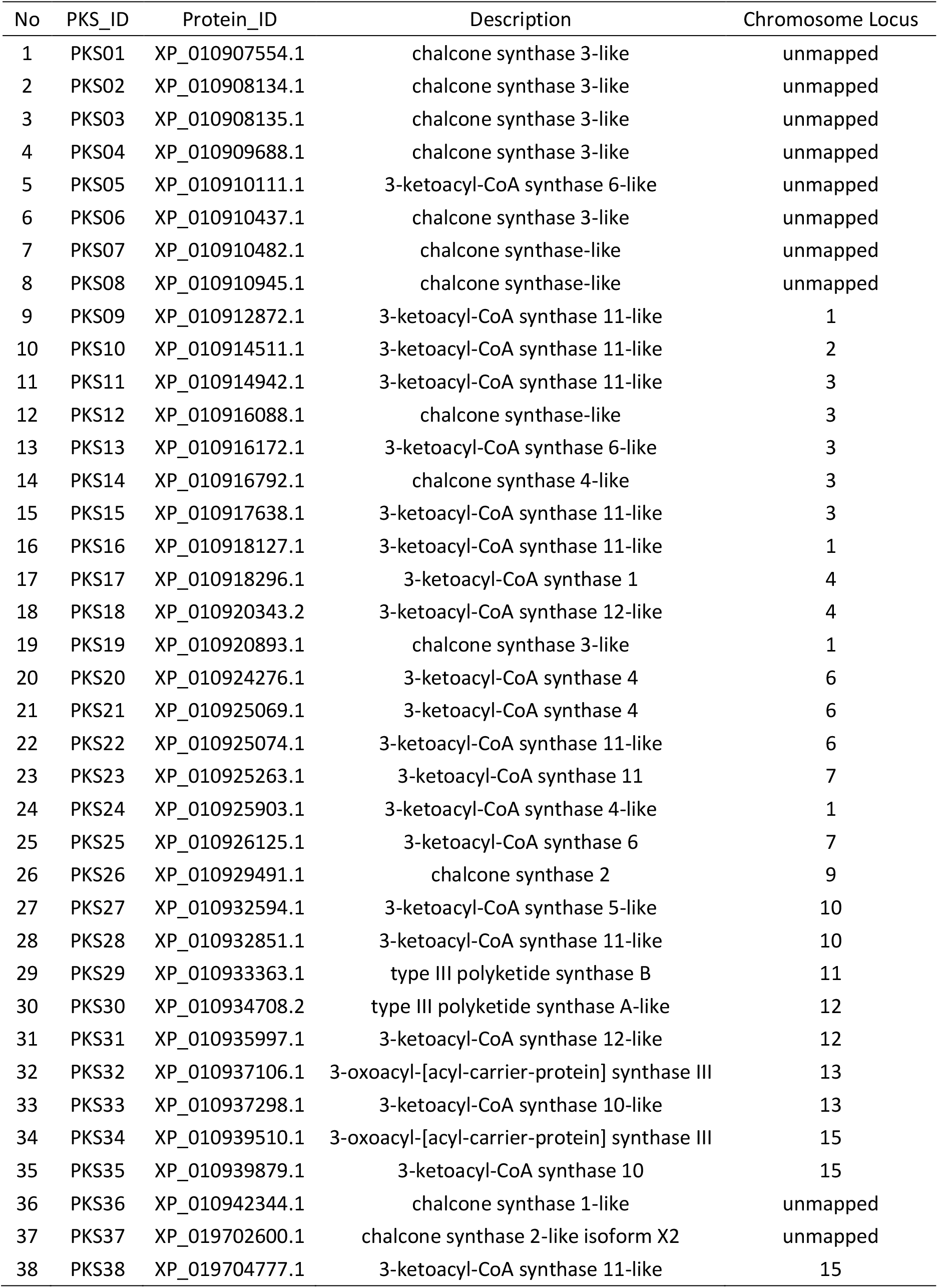
List of type III PKSs identified in *E. guineensis*.

| No | PKS_ID | Protein_ID | Description | Chromosome Locus |
| --- | --- | --- | --- | --- |
| 1 | PKS01 | XP_010907554.1 | chalcone synthase 3-like | unmapped |
| 2 | PKS02 | XP_010908134.1 | chalcone synthase 3-like | unmapped |
| 3 | PKS03 | XP_010908135.1 | chalcone synthase 3-like | unmapped |
| 4 | PKS04 | XP_010909688.1 | chalcone synthase 3-like | unmapped |
| 5 | PKS05 | XP_010910111.1 | 3-ketoacyl-CoA synthase 6-like | unmapped |
| 6 | PKS06 | XP_010910437.1 | chalcone synthase 3-like | unmapped |
| 7 | PKS07 | XP_010910482.1 | chalcone synthase-like | unmapped |
| 8 | PKS08 | XP_010910945.1 | chalcone synthase-like | unmapped |
| 9 | PKS09 | XP_010912872.1 | 3-ketoacyl-CoA synthase 11-like | 1 |
| 10 | PKS10 | XP_010914511.1 | 3-ketoacyl-CoA synthase 11-like | 2 |
| 11 | PKS11 | XP_010914942.1 | 3-ketoacyl-CoA synthase 11-like | 3 |
| 12 | PKS12 | XP_010916088.1 | chalcone synthase-like | 3 |
| 13 | PKS13 | XP_010916172.1 | 3-ketoacyl-CoA synthase 6-like | 3 |
| 14 | PKS14 | XP_010916792.1 | chalcone synthase 4-like | 3 |
| 15 | PKS15 | XP_010917638.1 | 3-ketoacyl-CoA synthase 11-like | 3 |
| 16 | PKS16 | XP_010918127.1 | 3-ketoacyl-CoA synthase 11-like | 1 |
| 17 | PKS17 | XP_010918296.1 | 3-ketoacyl-CoA synthase 1 | 4 |
| 18 | PKS18 | XP_010920343.2 | 3-ketoacyl-CoA synthase 12-like | 4 |
| 19 | PKS19 | XP_010920893.1 | chalcone synthase 3-like | 1 |
| 20 | PKS20 | XP_010924276.1 | 3-ketoacyl-CoA synthase 4 | 6 |
| 21 | PKS21 | XP_010925069.1 | 3-ketoacyl-CoA synthase 4 | 6 |
| 22 | PKS22 | XP_010925074.1 | 3-ketoacyl-CoA synthase 11-like | 6 |
| 23 | PKS23 | XP_010925263.1 | 3-ketoacyl-CoA synthase 11 | 7 |
| 24 | PKS24 | XP_010925903.1 | 3-ketoacyl-CoA synthase 4-like | 1 |
| 25 | PKS25 | XP_010926125.1 | 3-ketoacyl-CoA synthase 6 | 7 |
| 26 | PKS26 | XP_010929491.1 | chalcone synthase 2 | 9 |
| 27 | PKS27 | XP_010932594.1 | 3-ketoacyl-CoA synthase 5-like | 10 |
| 28 | PKS28 | XP_010932851.1 | 3-ketoacyl-CoA synthase 11-like | 10 |
| 29 | PKS29 | XP_010933363.1 | type III polyketide synthase B | 11 |
| 30 | PKS30 | XP_010934708.2 | type III polyketide synthase A-like | 12 |
| 31 | PKS31 | XP_010935997.1 | 3-ketoacyl-CoA synthase 12-like | 12 |
| 32 | PKS32 | XP_010937106.1 | 3-oxoacyl-[acyl-carrier-protein] synthase III | 13 |
| 33 | PKS33 | XP_010937298.1 | 3-ketoacyl-CoA synthase 10-like | 13 |
| 34 | PKS34 | XP_010939510.1 | 3-oxoacyl-[acyl-carrier-protein] synthase III | 15 |
| 35 | PKS35 | XP_010939879.1 | 3-ketoacyl-CoA synthase 10 | 15 |
| 36 | PKS36 | XP_010942344.1 | chalcone synthase 1-like | unmapped |
| 37 | PKS37 | XP_019702600.1 | chalcone synthase 2-like isoform X2 | unmapped |
| 38 | PKS38 | XP_019704777.1 | 3-ketoacyl-CoA synthase 11-like | 15 |

The number in oil palm is the highest that has been reported among other annual crops such as date palm. Date palm has only 9 and *A. thaliana* has 5 PKSs based on chalcone synthases and chalcone synthases like-proteins that deposited in the NCBI database. All of these PKSs were localized intracellular in the cytoplasma to build a polyketide product as been reported by (Mallika *et al*. 2011). According to our knowledge, this a first report describing PKSs in oil palm.

### Phyllogeny tree of oil palm PKSs

A phylogeny tree was performed to analyse evolutionary relationships among all PKSs. In this analysis, three clades were constructed and termed as PKS-A, -B and –C **Figure 1**. Clade A PKSs were characterized by the presence of N- and C-terminal domains of chalcone and stilbene synthases. Clade B PKSs have FAE1/Type III polyketide synthase-like protein and 3-Oxoacyl-[acyl-carrier-protein (ACP)] synthase III C terminal domains. While clade C PKSs have 3-Oxoacyl-[acyl-carrier-protein (ACP)] synthase and 3-Oxoacyl-[acyl-carrier-protein (ACP)] synthase III C terminal domains **Figure 2**. Clade PKS-A was comprised of all *P. dactylifera, A. thaliana* and some *E. guineensis* PKSs. Within this clade, two major branches were formed. The first branch was comprised of mostly *P. dactylifera* and *E. guineensis* indicating their close relationship among those two species. The second branch was comprised of most *A. thaliana*, few *P. dactylifera* and *E. guineensis* PKSs suggesting their conserved protein sequences between those species. The majority of *E. guineensis* PKSs were clustered in the PKS-B clade which might denote their specific function in the species. Clade PKS-C exclusively consisted of two *E. guineensis* PKSs i.e PKS32 and PKS34. Both PKSs have GTCILFG motif in their amino acid sequences that were absent in all analyzed PKSs and probably occurred as an evolutionary result of *E. guineensis*. The motif formed a naringenin-like structure near to the ACP binding site that has an unknown function up to now **Figure 3**.

**Figure 1.**
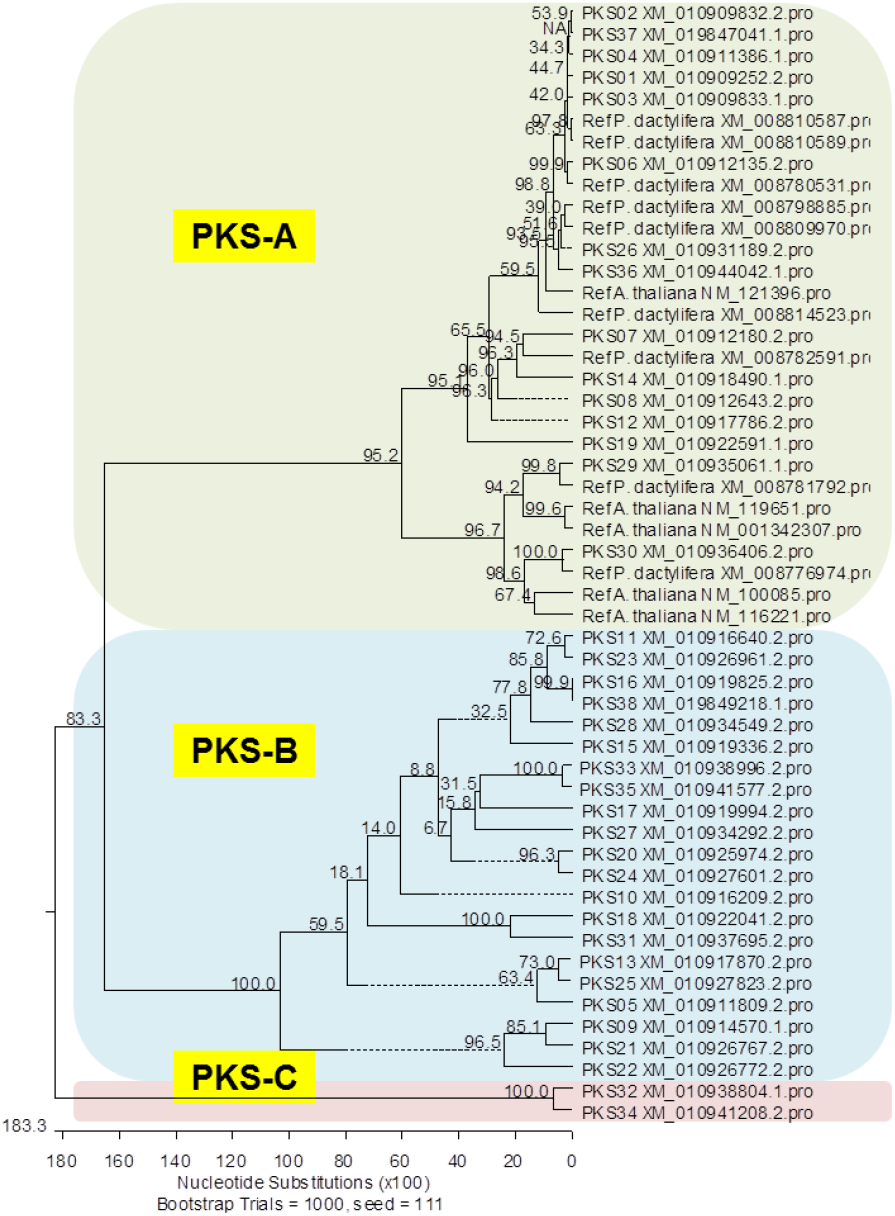
Molecular phylogeny of *E. guineensis* PKSs. The phylogram was gained through Neighbor-joining and described the relationship between amino acid sequences of PKSs extracted from 54 annotated proteins. Results from bootstrapping with 1,000 replicates were indicated.

**Figure 2.**
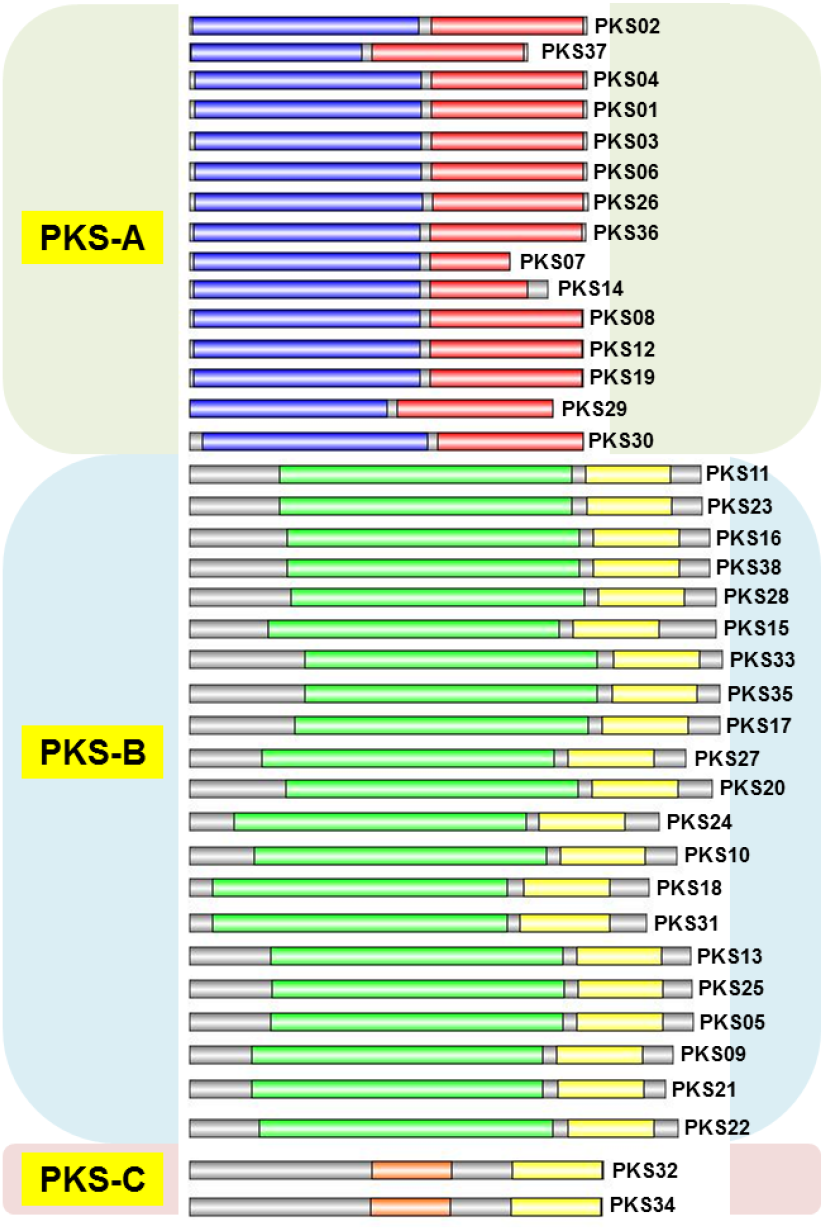
Molecular architecture of 38 *E. guineensis* PKS domains. Blue and Red indicated chalcone and stilbene synthases, N- and C-terminal domains, respectively. Green and yellow indicated FAE1/Type III polyketide synthase-like protein and 3-Oxoacyl-[acyl-carrier-protein (ACP)] synthase III C ter-minal domains, respectively. Orange indicated 3-Oxoacyl-[acyl-carrier-protein (ACP)] synthase domain.

**Figure 3.**
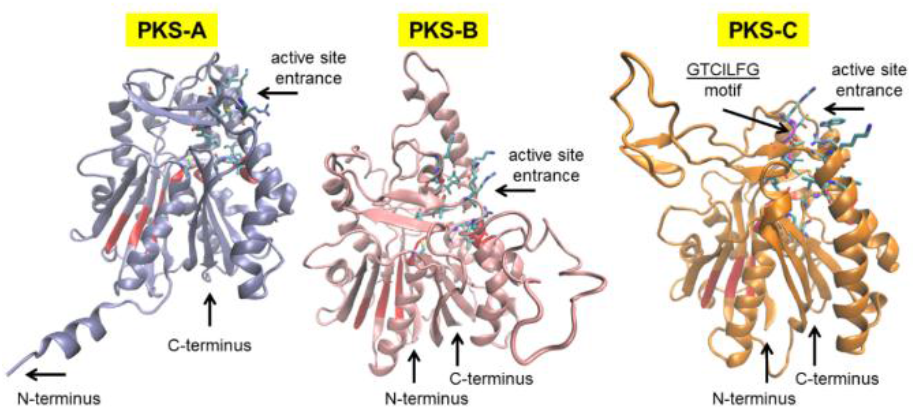
Comparison of 3-dimensional molecular structures of PKS02, PKS20 and PKS32. Each represented PKSA, -B, and -C clades, respectively. Red monomers indicated a conserved region for all PKS clades. PKS-C has a unique GTCILFG motif to form a naringenin-like structure which absent in other PKSs. N-, C-terminal end, and active site entrance are pointed by black arrows.

PKS in plants are specifically type III PKS, even though type III PKSs were also discovered in fungi and bacteria (Lim *et al*. 2016). Type III PKS is a homodimeric enzyme that performed a complete series of decarboxylation, condensation, and cylization reactions by utilizing CoA thioesters as substrates to tighten polyketide’s covalent attachment to the active site cysteine and does not entail the involvement of ACP domains as occurred in other type PKSs (Abe, 2008; Lussier *et al*. 2012; Tropf *et al*. 1995). The decarboxylation activates malonyl-CoA, iterative condensation couples the resulting acetyl anion to the growing ketide chain, and cylization forms the cyclized polyketide precursor of chalcone *via* an intramolecular Claisen condensation of the linear tetraketide intermediate (Austin and Noel, 2003).

The PKS clades illustrated the biodiversity and probably evolutionary story of the enzymes in *E. guineensis*. The first characterized type III PKS was the chalcone synthase of *Medicago sativa* (alfafa) to form chalcone that involved in the biosynthesis of flavonoid antimicrobial phytoalexins and anthocyanin pigments (Ferrer *et al*. 1999). The chalcone itself is an aromatic ketone that forms the central core for a variety of biological compounds (Schroeder, 1999). Its close-related enzyme, stilbene synthase, contribute to the resistance of woody tissues degradation and phytoalexins (Schroeder, 1997b). This heterogeneous lead to a large structural diversity and complex molecules that associated with divergent functions of the enzymes (Hertweck, 2009).

### Transcript level of 38 oil palm PKSs in response to *G. boninense* infection

Regarding the importance of *G. boninense* as the main fungal pathogen on oil palm in Indonesia, we analyzed whether any of these PKSs exhibited transcriptional up-regulation after *G. boninense* treatment. We performed artificial inoculation and conducted RNA sequencing to obtain the oil palm transcriptome profile. The transcriptional levels of all 38 PKSs were varied. The highest transcripts level was observed at PKS04 that expressed 117 folds compared to mock control. This suggested a role of this enzyme in producing a PK product that might needed as plant defense compound against *G. boninense*. Beside, other nine top up-regulated PKSs were shown by PKS01, -02, -37, -08, -03, -12, -07, -20, and PKS06 with 40.21, 29.49, 28.78, 12.38, 10.94, 7.71, 4.08, 2.07, and 1.90 folds, respectively **Table 2**. These PKSs could also involved in producing PK products for defense against *G. boninense*. Interestingly, all of these 10 PKSs shared high homology within Clade PKS-A, particularly PKS04, -01, -02, -37, and -03 as the top six up-regulated PKSs. Apparently, the chalcone and stilbene synthases, and N- and C-terminal domains in the PKSs, are essential to build PK products needed for defense.

**Table 2.** The transcription profile of all PKSs in response to *G. boninense* infection.

| PKS_ID | Treated (tpm) | Control (tpm) | Relative expression (FC) |
| --- | --- | --- | --- |
| PKS01 | 91.59 | 2.28 | 40.21 |
| PKS02 | 73.70 | 2.50 | 29.49 |
| PKS03 | 151.93 | 13.88 | 10.94 |
| PKS04 | 303.70 | 2.58 | 117.78 |
| PKS05 | 0.06 | 0.00 | #DIV/0! |
| PKS06 | 40.79 | 21.52 | 1.90 |
| PKS07 | 3.80 | 0.93 | 4.08 |
| PKS08 | 951.06 | 76.83 | 12.38 |
| PKS09 | 0.08 | 0.00 | #DIV/0! |
| PKS10 | 0.00 | 0.00 | #DIV/0! |
| PKS11 | 0.76 | 19.88 | 0.04 |
| PKS12 | 541.35 | 70.25 | 7.71 |
| PKS13 | 2.77 | 11.93 | 0.23 |
| PKS14 | 0.33 | 0.00 | #DIV/0! |
| PKS15 | 0.00 | 0.00 | #DIV/0! |
| PKS16 | 0.02 | 0.35 | 0.05 |
| PKS17 | 6.78 | 10.75 | 0.63 |
| PKS18 | 0.00 | 0.00 | #DIV/0! |
| PKS19 | 0.11 | 1.07 | 0.10 |
| PKS20 | 134.88 | 65.30 | 2.07 |
| PKS21 | 0.00 | 0.00 | #DIV/0! |
| PKS22 | 11.33 | 6.69 | 1.69 |
| PKS23 | 46.59 | 27.15 | 1.72 |
| PKS24 | 17.08 | 16.10 | 1.06 |
| PKS25 | 16.23 | 20.93 | 0.78 |
| PKS26 | 6.22 | 84.03 | 0.07 |
| PKS27 | 19.56 | 32.84 | 0.60 |
| PKS28 | 11.22 | 12.79 | 0.88 |
| PKS29 | 0.00 | 0.00 | #DIV/0! |
| PKS30 | 0.00 | 0.00 | #DIV/0! |
| PKS31 | 0.02 | 1.25 | 0.02 |
| PKS32 | 1.28 | 4.00 | 0.32 |
| PKS33 | 0.63 | 1.49 | 0.43 |
| PKS34 | 34.73 | 49.60 | 0.70 |
| PKS35 | 5.31 | 5.42 | 0.98 |
| PKS36 | 0.58 | 75.65 | 0.01 |
| PKS37 | 58.65 | 2.04 | 28.79 |
| PKS38 | 0.00 | 0.40 | 0.00 |

Polyketide secondary metabolite products exhibit divergent beneficial functions such as antibiotics, anticancer, immunosuppressive, antiparasitic, and antifungal (Staunton and Weissman, 2001; Wink, 2010). Nonetheless, most of the plant PKSs were characterized for their pharmaceutical properties since human health is considered more important rather than other properties such as phytoalexin (Dibyendu, 2015). Some plant metabolic compounds that characterized as phytoalexin and important as plant defense compound were reported in *A. thaliana*, *Zea mays*, *Lotus japonicus*, *Sorghum bicolor*, *Solanum lycopersicon*, *Oryza sativa, Papaver somniferum, Avena* spp. (Nuetzmann and Osbourn, 2014). These species are categorized as seasonal plants, and none was analyzed for annual crops such as *E. guineensis*. This study provides the first PKS identification on *E. guineensis* and the possible role of the biosynthesized products as plant defense biochemical compounds against *G. boninense*, but surely additional studies are needed to deliver more evidences.

## CONCLUSION

In conclusion, PK is one of a major secondary metabolite class found in many organisms. It has many beneficial functions such as antiparasitic phytoalexin in plants against pathogen invasion. Plant specific type III PKS is known using CoA thioesters as substrates rather than ACP domains as commonly used in other types PKS. This study revealed that *E. guineensis* have 38 PKSs that scattered in the chromosome. These PKSs were clustered into three clades illustrating their molecular relationship based on amino acid sequences. PKS04 encoding gene was the highest upregulated gene in response to *G. boninense* infection. It indicated its contribution in the biosynthesis of a PK product that might needed as plant defense compounds to inhibit pathogen development. A further study such as gene cluster knock out could be done to characterize the gene cluster function. According to our knowledge this is a first report revealing all PKSs in oil palm and predict its role in plant defense.

## ACKNOWLEDGEMENTS

We thank the top management of PT SMART Tbk. who provided the financial support for the project under a research code 3.3.1.031. We also thank Corry Korita Neryceka and Widyartini who provide assistance in the administration process of publishing this article.

## AUTHORS’ NOTE

The author(s) declare(s) that there is no conflict of interest regarding the publication of this article. Authors confirmed that the data and the paper are free of plagiarism.

